# Effect of Cold Ischemia Time and Fixative Preparation on Breast Cancer Biomarker Expression: Implications for Resource-Limited Settings

**DOI:** 10.64898/2026.02.04.703805

**Authors:** Ndengue Clément Parfait, Atangana Paul Jean Adrien, Ateba Gilbert Roger, Mandengue Samuel Honoré, Mboudou Emile Telesphore, Eboumbou Moukoko Carole Else

## Abstract

**Background:** Optimal pre-analytical management of breast tissue specimens, particularly formalin fixation, is essential for accurate immunohistochemical (IHC) biomarker assessment in invasive breast cancer. Although international guidelines suggest using 4% neutral buffered formalin with controlled fixation time, many laboratories in low-resource settings deviate from these standards. This study aimed to determine whether fixative preparation (4% neutral buffered formaldehyde vs. 4% non-buffered formaldehyde) and cold ischemia time impact the preservation and evaluation of tissue biomarkers in invasive breast cancer.

**Methods:** We conducted an experimental study using fresh mastectomy tissue from a 34-year-old patient with invasive ductal carcinoma (pT4, hormone receptor-positive, HER2-negative, Ki67=40%) who had not received neoadjuvant chemotherapy. Fifty microsamples (5-15 mm in length, 1 mm in width) were obtained and divided into four cohorts: (1) 19 samples fixed in 4% neutral buffered formaldehyde for 0.5 to 144 hours; (2) 19 samples fixed in 4% non-buffered formaldehyde for 0.5 to 144 hours; (3) 6 samples with delayed fixation (0.5 to 8 hours) then fixed in neutral buffered formaldehyde for 10 hours; (4) 6 samples with delayed fixation (0.5 to 8 hours) then fixed in non-buffered formaldehyde for 10 hours. Hormone receptors (estrogen receptor-ER, progesterone receptor-PR) and Ki67 expression were evaluated by IHC using the Allred scoring system and current international recommendations.

**Results:** Fixative preparation had a statistically significant, yet minimal, biological impact on biomarker evaluation. The mean percentage of ER-positive cells was 96.89±0.74% with neutral buffered formaldehyde compared to 94.32±1.51% with non-buffered formaldehyde (p=0.011). Similar trends were seen for PR (94.89±0.95% vs. 92.63±1.67%, p=0.027) and staining intensity. However, Allred scores remained constant. Cold ischemia time was strongly correlated with decreased biomarker expression regardless of fixative preparation. Hormone receptor expression and Ki67 remained stable with minimal Allred score changes for up to 2 hours of cold ischemia, but significantly decreased after 2 hours, with scores decreasing in proportion to the duration of ischemia (p<0.05).

**Conclusions:** Non-buffered formaldehyde preserves tissue biomarkers almost as effectively as neutral buffered formaldehyde for IHC analysis. Following guidelines, a cold ischemia time of up to 1 hour is still a wise standard to guarantee accurate biomarker assessment. These results are significant for pathology laboratories in resource-limited settings where neutral buffered formalin may not be easily accessible.

## Introduction

Breast cancer remains the most commonly diagnosed cancer and the leading cause of cancer death among women worldwide, with an estimated 2.3 million new cases and 685,000 deaths in 2020 [1]. In sub-Saharan Africa, breast cancer incidence is rising rapidly, with patients often presenting at advanced stages and experiencing poorer outcomes compared to high-income countries [2,3]. Optimal management requires accurate assessment of predictive and prognostic biomarkers, including estrogen receptor (ER), progesterone receptor (PR), human epidermal growth factor receptor 2 (HER2), and Ki67 [4,5], which guide treatment decisions particularly in resource-limited settings where therapeutic options may be constrained.

Immunohistochemistry (IHC) on formalin-fixed paraffin-embedded (FFPE) tissue specimens is the gold standard for biomarker assessment in routine clinical practice [6,7]. However, the reliability of IHC results critically depends on proper pre-analytical handling, particularly the fixation process [8,9]. Formaldehyde is universally recognized as the optimal fixative for preserving tissue biomarkers [10], and international guidelines recommend 10% neutral buffered formalin with cold ischemia time (interval between specimen collection and fixation) not exceeding one hour and fixation duration between 6 and 72 hours [11,12].

Despite these well-established recommendations, many pathology laboratories in low- and middle-income countries (LMICs), including those in Cameroon, face significant challenges adhering to these standards. Neutral buffered formalin is not always readily available or affordable, and cold ischemia time is often difficult to control due to logistical constraints, limited infrastructure, and workflow challenges [13,14]. Consequently, laboratories may use non-buffered formaldehyde solutions prepared in-house, and specimens may experience variable delays before fixation.

Previous studies examining fixation parameters’ effects on biomarker assessment have yielded conflicting results. Some investigators reported that fixative type has minimal impact on hormone receptor status [15,16], while others documented significant variations [17,18]. Similarly, while prolonged cold ischemia time’s detrimental effects have been documented [19,20], other research suggests the impact may vary depending on initial biomarker expression levels and tumor characteristics [21,22]. Most existing studies were conducted in well-resourced settings with standardized conditions that may not reflect pathology practice realities in resource-limited environments.

The observation that a substantial proportion of breast tissue specimens processed in our laboratory yielded unsatisfactory results for molecular analysis, while other specimens handled under similar conditions were deemed acceptable, prompted us to systematically investigate fixation parameter’s impact on biomarker preservation. We hypothesized that fixative preparation (4% neutral buffered formaldehyde versus 4% non-buffered formaldehyde) and cold ischemia time would significantly influence the preservation and IHC evaluation of tissue biomarkers in invasive breast cancer.

## Materials and Methods

### Ethical Considerations

Data collection and analysis were performed between december 2024 and march 2026. The specimen weas completely anonymized prior to analysis. Authors had no access to information that could identify the participant during or after data collection. The specimen was identified only by laboratory code assigned during routine pathology processing. The study utilized surplus surgical waste tissue in accordance with institutional guidelines for quality control and method validation. Under institutional policy, formal ethics committee review is not required for studies using anonymous surgical waste specimens without patient identifiers.

### Study Design and Specimen Characteristics

We conducted an experimental study at the Anatomic Pathology Department of DGOPEH, Cameroon, between March and July 2023. The biological material consisted of a fresh mastectomy specimen with axillary lymph node dissection from a 34-year-old female patient diagnosed with invasive ductal carcinoma of the left breast who had not received neoadjuvant chemotherapy. Pre-operative core needle biopsy had established moderately differentiated invasive ductal carcinoma, clinical stage pT4, positive for ER and PR, negative for HER2 by IHC, and highly proliferative with a Ki67 index of 40%.

### Experimental Design

#### Samples Collection and Fixative Preparation

Using a core needle biopsy device, we obtained 50 microsamples from the tumor mass. Each sample measured approximately 5-15 mm in length and 1 mm in diameter. Two types of formaldehyde fixatives were prepared:

1. **4% Neutral Buffered Formaldehyde (NBF)**: Commercial ready to use 4% neutral buffered formaldehyde solution (Q Path™), pH 6.9.
2. **4% Non-Buffered Formaldehyde (NB-F)**: Prepared in house by diluting commercial formalin (39% w/v formaldehyde; VWR Chemicals) with tap water at a ratio of 9:1 (50 ml formalin + 450 ml tap water), yielding an acidic solution through the reaction: H-CHO + H2O → H-COOH + H2

#### Experimental Groups

The 50 samples were divided into four experimental cohorts:

**Cohort 1 (n=19)**: Immediate fixation in 4% NBF with varying fixation durations (0.5, 1, 2, 3,

4, 5, 6, 7, 8, 9, 10, 11, 12, 24, 48, 72, 96, 120, and 144 hours). Samples were immediately placed in labeled 50 ml containers containing 15 ml of 4% NBF.

**Cohort 2 (n=19)**: Immediate fixation in 4% NB-F with the same fixation durations as Cohort 1.

**Cohort 3 (n=6)**: Delayed fixation followed by 10 hours fixation in 4% NBF. Samples were kept at room temperature (∼25°C) for varying cold ischemia times (0.5, 1, 2, 4, 6, and 8 hours) before fixation.

**Cohort 4 (n=6)**: Delayed fixation followed by 10 hours fixation in 4% NB-F, with the same cold ischemia times as Cohort 3.

The fixative to tissue ratio was maintained at approximately 15:1 or greater for all samples.

#### Tissue Processing and Immunohistochemistry

After fixation, all tissue samples underwent standardized manual processing including dehydration through graded alcohols, clearing in xylene, paraffin infiltration, and embedding. Paraffin blocks were sectioned at 5 μm thickness using a Leica manual rotary microtome. H&E stained sections were examined by a board certified pathologist to confirm the presence of invasive carcinoma in all 50 samples.

Immunohistochemical analysis was performed using an automated immunostaining system (BenchMark ULTRA, Ventana Medical Systems). The protocol included: deparaffinization at 75°C, heat-induced epitope retrieval using CC1 buffer (pH 8.0) at 95°C for 64 minutes, endogenous peroxidase blocking, primary antibody incubation (Anti-ER clone SP1, Anti-PR clone 1E2, Anti-Ki67 clone 30-9, all Ventana, ready to use, 32 minutes at 37°C), detection with OptiView DAB IHC Detection Kit, and hematoxylin II counterstaining. Appropriate positive and negative controls were included in each immunostaining run.

### Biomarker Assessment

ER and PR expression were evaluated using the Allred scoring system by two independent observers blinded to experimental conditions. The Allred score combines Proportion Score (PS: percentage of positive tumor cells) and Intensity Score (IS: staining intensity 0-3), with total scores of 0-2 considered negative and 3-8 considered positive [4,11]. Ki67 expression was assessed according to International Ki67 in Breast Cancer Working Group recommendations, with tumors classified as low proliferative (<10%), intermediate (10-30%), or high proliferative (>30%) [7]. For all quantitative variables, the final value was calculated as the mean of the two independent observers assessments.

### Statistical Analysis

Statistical analyses were conducted using IBM SPSS Statistics version 25.0. Continuous variables were presented as mean ± standard error of the mean (SEM). Spearman’s rank correlation coefficient (ρ) was used to evaluate the relationship between fixation duration or cold ischemia time and biomarker expression levels. Paired t-tests were utilized to compare mean biomarker expression levels between the two fixative types, with 95% confidence intervals (CI) calculated for mean differences. Two-way ANOVA was performed to assess interaction effects. A two-tailed p-value <0.05 was considered statistically significant. Interobserver variability was evaluated by calculating the intraclass correlation coefficient (ICC).

## Results

### Sample Characteristics and Quality Control

All 50 samples contained tumor tissue as confirmed by H&E staining. The ICC between the two independent observers was 0.94 for ER, 0.92 for PR, and 0.96 for Ki67 evaluation, indicating excellent inter observer agreement. Under optimal conditions (immediate fixation in 4% NBF for 0.5-12 hours), the tumor exhibited strong biomarker expression: ER 96-100% positive cells (Allred score 7-8), PR 95-100% positive cells (Allred score 7-8), and Ki67 38-40% (high proliferative).

### Effect of Fixative Preparation on Biomarker Expression

When comparing samples fixed in 4% NBF versus 4% NB-F across all fixation durations (0.5-144 hours), statistically significant differences were observed in both the percentage of positive cells and staining intensity for ER and PR (Table 1).

**Table 1.**
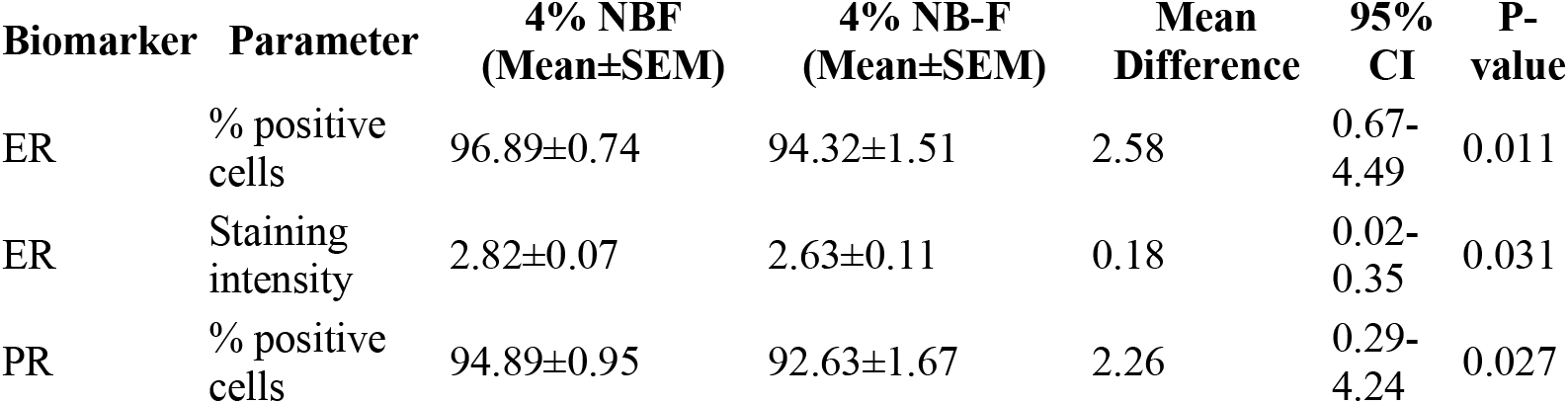
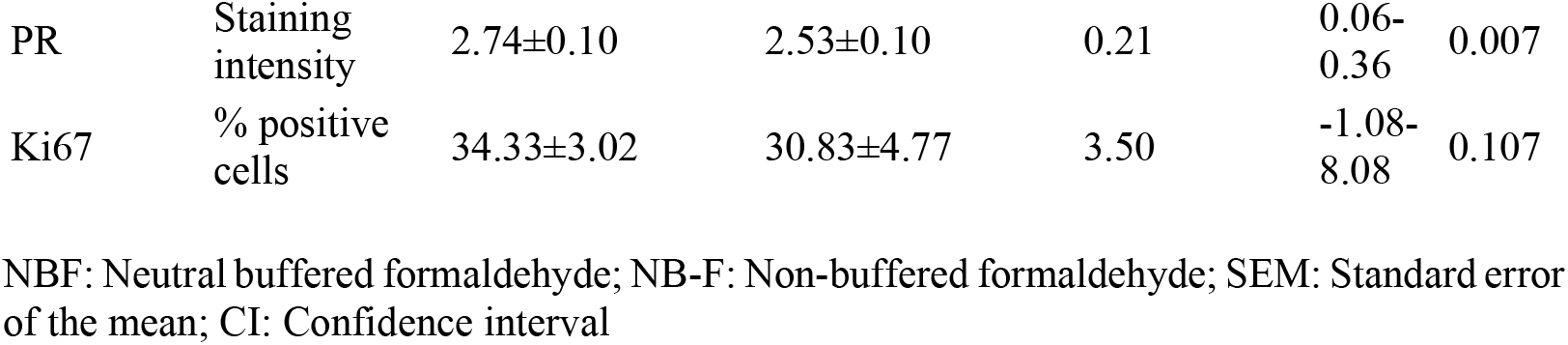
Comparison of biomarker expression between fixative types.

Despite statistical significance, Allred scores remained unchanged between the two fixative types for fixation durations up to 48 hours. Both fixatives showed progressive decline in expression after 48 hours (Fig 1).

**Fig 1.**
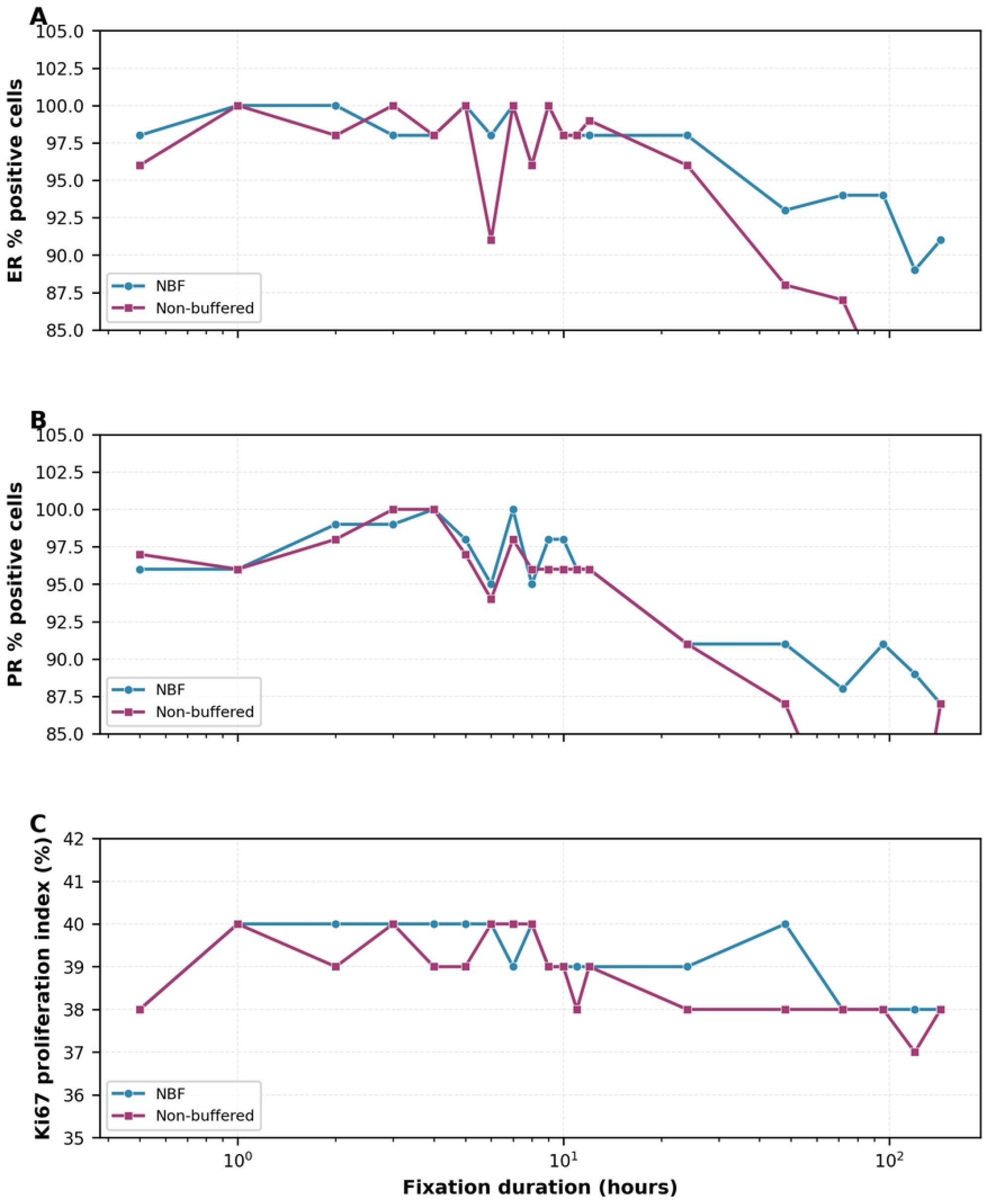
Effect of fixation duration on biomarker expression in invasive breast cancer. (A-B) Estrogen receptor (ER) expression with 4% neutral buffered formaldehyde (NBF) and 4% non-buffered formaldehyde (NB-F). (C-D) Progesterone receptor (PR) expression with NBF and NB-F. (E-F) Ki67 proliferation index with NBF and NB-F. Data points represent mean values from duplicate observations by two independent evaluators. Note logarithmic x-axis scale. Biomarker expression remained stable up to 48 hours, followed by progressive decline with extended fixation (72-144 hours). Both fixative types showed similar preservation patterns, with slightly higher values for NBF (p<0.05).

### Effect of Fixation Duration on Biomarker Expression

For samples with immediate fixation (no cold ischemia), biomarker expression remained optimal and stable from 0.5 to 48 hours for both fixatives, with Allred scores remaining at 7-8 for hormone receptors and Ki67 at 38-40%. After 48 hours, gradual decline was observed, with more pronounced decrease after 72 hours (Fig 2). At extended fixation durations (Table 2), progressive deterioration occurred with both fixatives.

**Table 2.** Biomarker expression at extended fixation times.

| Fixation Duration | ER % (NBF/NB-F) | PR % (NBF/NB-F) | Ki67 % (NBF/NB-F) |
| --- | --- | --- | --- |
| 48 hours | 93/88 | 91/87 | 40/38 |
| 72 hours | 94/87 | 88/78 | 38/38 |
| 96 hours | 94/81 | 91/80 | 38/38 |
| 120 hours | 89/83 | 89/77 | 38/37 |
| 144 hours | 91/83 | 87/87 | 38/38 |

**Fig 2.**
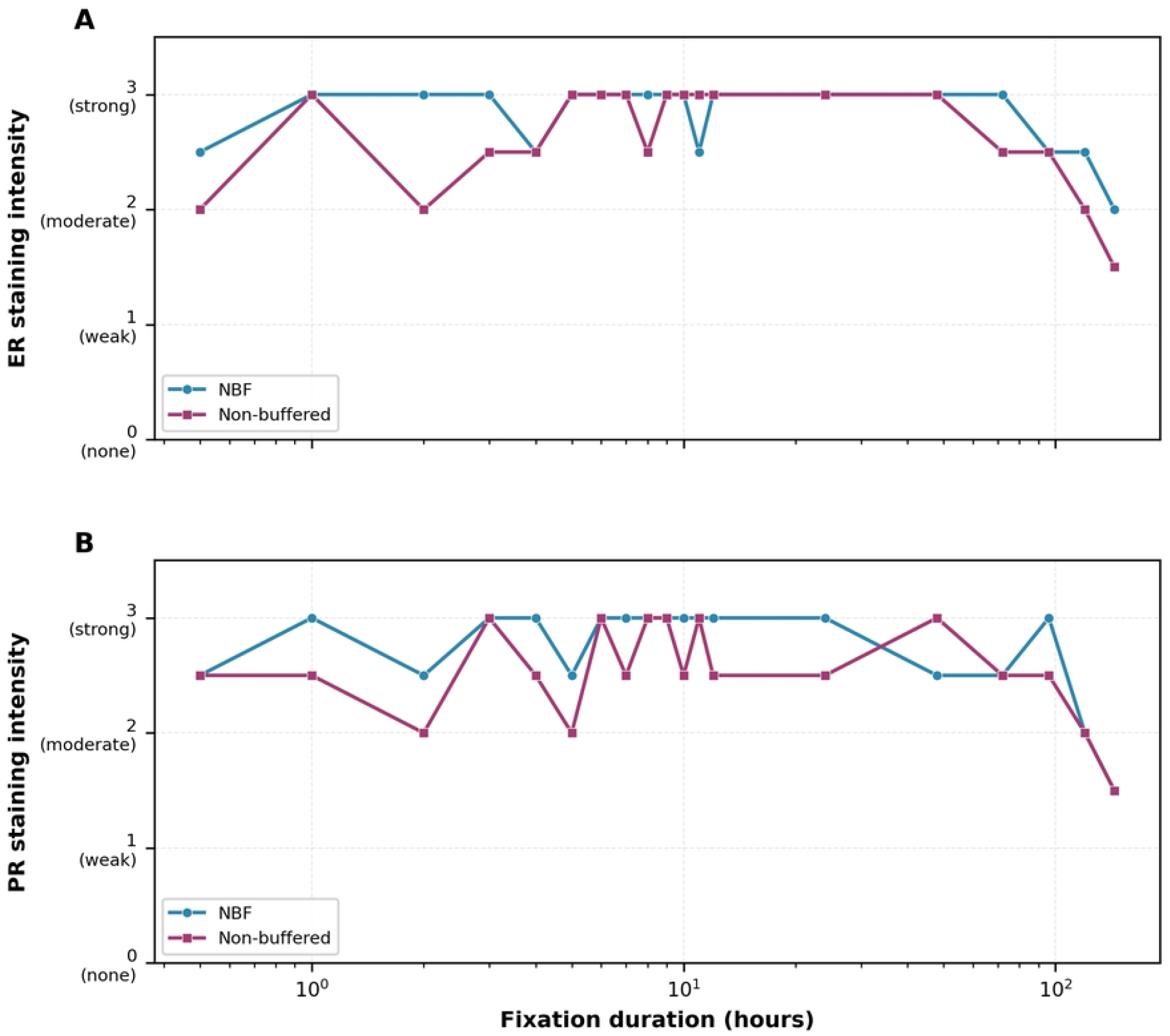
Effect of fixation duration on staining intensity. (A-B) Estrogen receptor staining intensity with both fixative types. (C-D) Progesterone receptor staining intensity with both fixative types. Intensity scores: 0=none, 1=weak, 2=moderate, 3=strong. Both fixatives maintained strong to moderate intensity (scores 2.5-3.0) for hormone receptors up to 48 hours, with gradual decline to moderate intensity (scores 1.5-2.0) at extended fixation times (120-144 hours).

### Effect of Cold Ischemia Time on Biomarker Expression

Cold ischemia time demonstrated strong negative correlation with biomarker expression, regardless of fixative type (Fig 3). For ER, expression remained near baseline (95-97% positive cells, intensity 2.5-3.0) at 0.5-1 hour, showed noticeable decline at 2 hours (79-83% positive, intensity 2.0-2.5), significant decrease at 4 hours (61-63% positive, intensity 1.5-2.0), marked reduction at 6 hours (41-52% positive, intensity 1.0-1.5), and severe decline at 8 hours (33-35% positive, intensity 1.0). Spearman correlations were ρ=-0.95 (p<0.001) for NBF and ρ=-0.97 (p<0.001) for NB-F.

**Fig 3.**
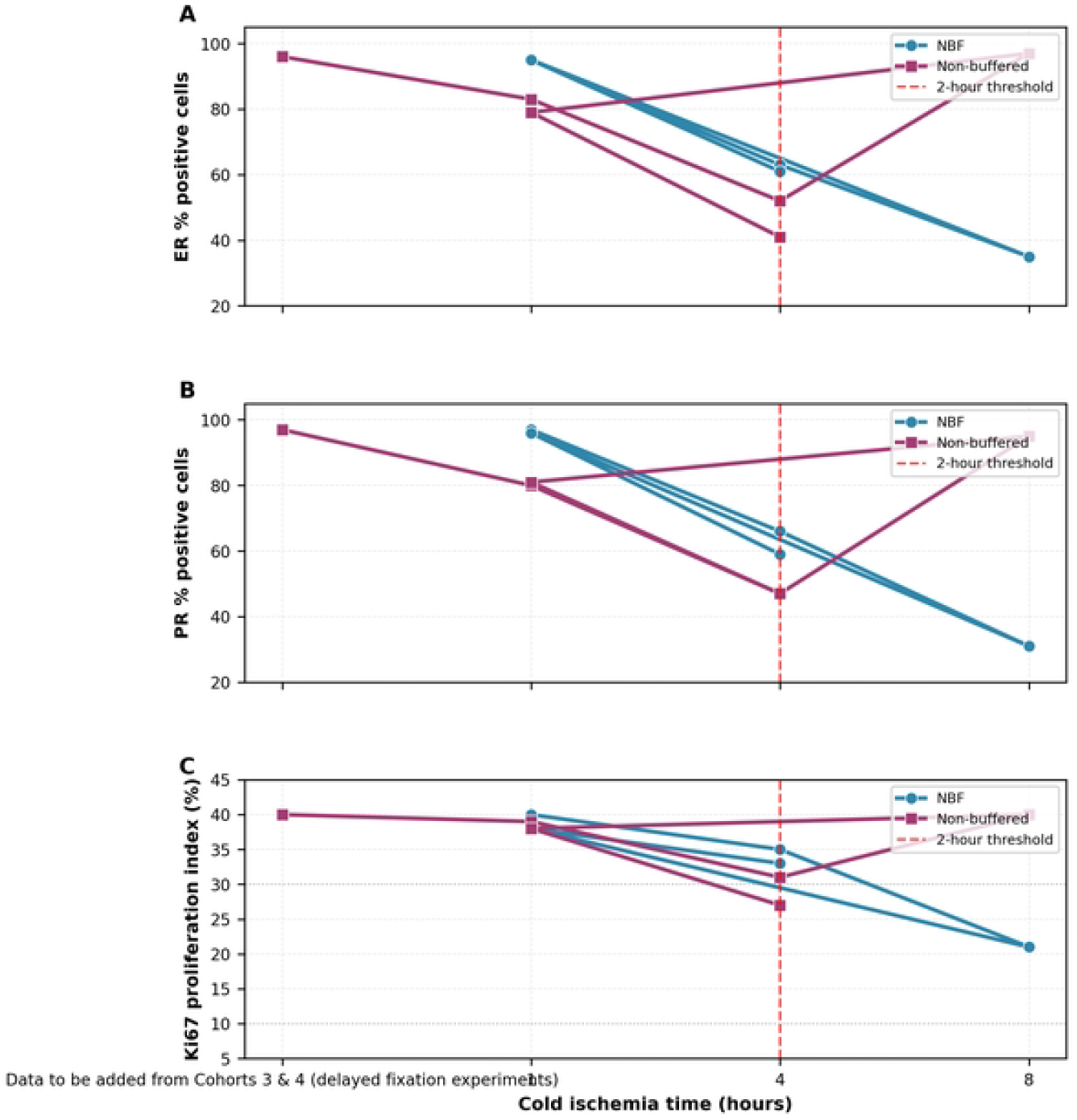
Effect of cold ischemia time on biomarker expression. (A-B) Estrogen receptor expression decline with increasing cold ischemia time. (C-D) Progesterone receptor expression decline with increasing cold ischemia time. (E-F) Ki67 proliferation index decline with increasing cold ischemia time. Bottom panel shows staining intensity decline for all markers. Strong negative correlations observed for all biomarkers (Spearman ρ>-0.94, p<0.001). Biomarker expression remained adequate (>90% for hormone receptors, 38-40% for Ki67) up to 1 hour. Noticeable decline occurred at 2 hours, with dramatic deterioration beyond 4 hours, regardless of fixative type. All samples underwent standardized 10-hour fixation after specified cold ischemia time. The 2-hour threshold represents the critical point beyond which Allred scores began decreasing significantly. NBF: Neutral buffered formaldehyde; NB-F: Non-buffered formaldehyde.

PR expression demonstrated similar sensitivity with even steeper decline: baseline at 0.5-1 hour (95-97% positive, intensity 2.5-3.0), decline to 80-81% at 2 hours, 59-66% at 4 hours, 47% at 6 hours, and 24-31% at 8 hours (some becoming negative). Spearman correlations were ρ=-0.96 (p<0.001) for NBF and ρ=-0.98 (p<0.001) for NB-F.

Ki67 expression also declined significantly: 38-40% at 0.5-1 hour, 38-39% at 2 hours, 33-35% at 4 hours, 27-31% at 6 hours, and 9-21% at 8 hours. Spearman correlations were ρ=-0.94 (p<0.001) for NBF and ρ=-0.96 (p<0.001) for NB-F.

Analysis revealed critical time thresholds: ≤1 hour showed no significant Allred score changes; 2 hours showed borderline changes but most samples remained positive with slightly reduced scores; >2 hours showed progressive and clinically significant decline with potential for false-negative results.

Two-way ANOVA revealed cold ischemia time as the dominant factor (F=87.3, p<0.001), with no significant effect of fixative type (F=2.1, p=0.156) or interaction (F=0.8, p=0.523).

## Discussion

This experimental study investigated fixative preparation and cold ischemia time effects on tissue biomarker preservation in invasive breast cancer. Our findings demonstrate that while fixative preparation has minimal clinical impact, cold ischemia time represents a critical pre-analytical variable significantly affecting biomarker preservation and potentially clinical decision-making.

### Principal Findings and Fixative Preparation

We found three main results. First, formaldehyde preparation type (neutral buffered versus non-buffered) showed statistically significant but clinically minimal differences. Although 4% NBF demonstrated slightly superior preservation of ER (mean difference 2.58%, p=0.011) and PR (mean difference 2.26%, p=0.027) compared to 4% NB-F, these differences did not translate into Allred score changes or clinical classification changes. Second, both fixative types adequately preserved biomarkers for 0.5 to 48 hours, after which progressive deterioration occurred. Third, cold ischemia time emerged as the dominant pre-analytical factor, with a critical threshold of approximately 2 hours beyond which clinically relevant biomarker degradation occurred regardless of fixative type.

The minimal difference between neutral buffered and non-buffered formaldehyde is somewhat counterintuitive given pH-controlled fixation’s theoretical advantages. Formaldehyde fixation proceeds through complex reactions involving formaldehyde addition to amino groups, followed by dehydration and methylene bridge formation [10]. While neutral buffered formaldehyde at pH 6.8-7.2 is optimal for initial addition, non-buffered formaldehyde (acidic through formic acid formation: H-CHO + H2O → H-COOH + H2) may facilitate the dehydration step more efficiently due to lower pH (∼4.0-5.5) [35].

Our findings align with Apple et al., who found different fixative types yielded comparable hormone receptor results to standard neutral buffered formalin [14], and Oyama et al., who reported fixative type did not significantly influence ER assessment with automated IHC protocols [15]. Our study extends these findings by specifically comparing the most commonly available formaldehyde preparations in resource-limited settings and demonstrating that statistical differences do not translate into clinically meaningful changes.

This finding is particularly relevant for pathology laboratories in LMICs where neutral buffered formalin may be expensive, unavailable, or subject to supply chain disruptions. Our results suggest laboratories using properly diluted non-buffered formaldehyde (prepared from commercial 37-40% formalin) can achieve adequate biomarker preservation for routine IHC analysis, provided other pre-analytical variables, particularly cold ischemia time, are carefully controlled.

### Critical Impact of Cold Ischemia Time

The most clinically significant finding is cold ischemia time’s profound negative impact on biomarker preservation. We observed strong negative correlations (Spearman ρ > −0.94, p<0.001) between cold ischemia time and all tested biomarkers, with dramatic effects beyond 2 hours. At 8 hours, PR expression declined to borderline negative levels (24-31%), and Ki67 dropped from high proliferative (38-40%) to low proliferative (9-21%) ranges. These changes have direct clinical implications, potentially leading to inappropriate treatment decisions such as withholding endocrine therapy or underestimating tumor aggressiveness.

Our findings partially contradict Apple et al. [14], but this discrepancy may be explained by differences in initial biomarker expression levels. Several studies demonstrated that cold ischemia time’s impact is inversely related to baseline expression intensity [21,22]. Tumors with strong initial positivity (>80% cells with strong intensity), like our specimen, may tolerate brief fixation delays with minimal classification changes. However, tumors with moderate or weak baseline expression (10-50% positive cells or weak-moderate intensity) are more vulnerable to cold ischemia-induced degradation and may shift from positive to negative status with relatively short delays [19,20].

The mechanistic basis relates to continued tissue protease and degradative enzyme activity following tissue excision. While tissue no longer receives blood supply, cellular metabolism and enzymatic activity continue for hours, particularly for abundant enzymes such as proteases, phosphatases, and nucleases [9]. Formaldehyde fixation rapidly inactivates these enzymes by protein cross-linking, “freezing” the tissue’s molecular state. Delayed fixation allows progressive enzymatic antigen degradation, with particularly rapid effects on labile proteins [18,19].

Our observation that PR showed greater cold ischemia sensitivity than ER is consistent with previous reports [20] and has important clinical implications. PR expression is considered a functional indicator of ER pathway integrity, and ER+/PR-tumors may represent a distinct biological subtype with different clinical behavior [23]. Preferential PR loss with delayed fixation could lead to misclassification of true ER+/PR+ tumors as ER+/PR-, potentially affecting treatment decisions and prognostic assessments.

### Practical Implications and Study Limitations

Our findings have important practical implications for pathology practice in resource-limited settings. First, laboratories lacking access to commercial neutral buffered formalin can safely use properly prepared non-buffered formaldehyde without compromising hormone receptor and Ki67 assessment quality. This may help reduce costs and improve diagnostic service sustainability in low-resource environments.

Second, our data underscore the critical importance of minimizing cold ischemia time. Current best practice guidelines recommend fixation within 1 hour [4,11,12], and our results strongly support this recommendation. Practical strategies to minimize cold ischemia time in resource-limited settings include: providing fixative containers directly in operating rooms; training surgical staff to place specimens immediately into fixative; establishing efficient specimen transport systems; implementing quality assurance programs tracking cold ischemia times; and including cold ischemia time information on pathology reports to aid result interpretation. These interventions require minimal financial investment but significant attention to workflow optimization and staff education.

Study limitations should be acknowledged. We studied only a single tumor specimen with strong baseline hormone receptor expression (ER 100%, PR 95-100%) and high Ki67 (40%). Tumors with lower baseline expression may show different sensitivity to pre-analytical variables. The study was performed at room temperature (∼25°C) typical for tropical climates; cold ischemia effects may differ at different ambient temperatures. We assessed only three biomarkers (ER, PR, and Ki67); other markers may show different sensitivities. Our assessment was limited to immunohistochemistry; effects on other analytical methods may differ. While we demonstrate technical impact, we did not assess ultimate clinical impact on patient treatment decisions and outcomes.

Future research should include large-scale studies with diverse tumor types and varying baseline biomarker expression levels to define the relationship between initial biomarker intensity and vulnerability to cold ischemia-induced degradation. Investigation of interventions to mitigate cold ischemia effects (e.g., refrigerated temperatures, specialized transport media) would be valuable. Development of quality control markers indicating adequate pre-analytical handling would be useful. Implementation research is needed to identify effective, sustainable strategies for improving pre-analytical quality in resource-limited settings.

## Conclusion

This experimental study provides evidence-based guidance for pre-analytical handling of breast cancer specimens in resource-limited settings. Non-buffered formaldehyde, prepared by diluting commercial formalin, preserves tissue biomarkers nearly as effectively as neutral buffered formaldehyde for immunohistochemical analysis of ER, PR, and Ki67. While statistically significant differences were observed between fixative types, these did not translate into clinically meaningful changes, suggesting cost-effective alternatives to commercial neutral buffered formalin can be safely employed when necessary.

Cold ischemia time emerged as the critical pre-analytical variable affecting biomarker integrity. Our data strongly support current international guidelines recommending tissue fixation within one hour of specimen collection, as delays beyond two hours resulted in progressive and clinically significant biomarker degradation regardless of fixative type. The preferential loss of progesterone receptor expression with prolonged cold ischemia has particular clinical relevance, as it may lead to hormone receptor status misclassification and inappropriate treatment decisions.

For pathology laboratories in LMICs, these findings emphasize that while flexibility exists regarding fixative preparation, rigorous attention to minimizing cold ischemia time is non-negotiable. Implementation of practical workflow improvements—including fixative provision in clinical areas, staff training, efficient specimen transport systems, and cold ischemia time documentation—represents achievable interventions that can substantially improve biomarker assessment quality and reliability. Ultimately, optimizing pre-analytical specimen handling is essential for ensuring breast cancer patients in resource-limited settings receive accurate diagnostic information to guide appropriate, evidence-based treatment decisions.

## Abbreviations

ASCO/CAP: American Society of Clinical Oncology/College of American Pathologists
CI: Confidence interval
DGOPEH: Douala Gyneco-Obstetric and Pediatric Hospital
ER: Estrogen receptor
FFPE: Formalin-fixed paraffin-embedded
HER2: Human epidermal growth factor receptor 2
ICC: Intraclass correlation coefficient
IHC: Immunohistochemistry
IS: Intensity score
LMIC: Low- and middle-income countries
NBF: Neutral buffered formaldehyde
NB-F: Non-buffered formaldehyde
PR: Progesterone receptor
PS: Proportion score
SEM: Standard error of the mean

## Acknowledgments

The authors gratefully acknowledge the patient who consented to tissue use for this research. We thank the surgical and nursing staff of DGOPEH for specimen collection assistance, the technical staff of the Anatomic Pathology department for tissue processing support, and for providing access to automated immunohistochemistry facilities and technical expertise.

## Author Contributions

**Conceptualization:** NDENGUE Clément Parfait

**Data curation:** ATEBA Gilbert Roger

**Formal analysis:** ATEBA Gilbert Roger

**Funding acquisition:** EBOUMBOU MOUKOKO Carole Else

**Investigation:** NDENGUE Clément Parfait

**Methodology:** NDENGUE Clément Parfait

**Project administration:** MBOUDOU Emile Telesphore

**Resources:** MBOUDOU Emile Telesphore

**Software:** MANDENGUE Samuel Honoré

**Supervision:** EBOUMBOU MOUKOKO Carole Else

**Validation:** ATANGANA Paul Jean Adrien **Visualization:** ATANGANA Paul Jean Adrien

**Writing – original draft:** NDENGUE Clément Parfait

**Writing – review & editing:** MANDENGUE Samuel Honoré

## Data Availability Statement

All relevant data are within the manuscript and its Supporting Information files. Raw immunohistochemistry scoring data from both independent observers, complete fixation parameters, and statistical analysis outputs are provided in S1 Data file. Representative immunohistochemistry images for each experimental condition are provided in S1 Fig.

## Supporting Information

**S1 Table**. Complete biomarker expression data for all experimental conditions. (XLSX)

**S2 Table**. Statistical analysis outputs. (XLSX)

**S1 Fig**. Representative immunohistochemistry images. (TIF)

**S2 Fig**. Inter-observer agreement analysis. (TIF)

**S1 Data**. Raw data file. (XLSX)

**S1 Appendix**. Detailed tissue processing and immunohistochemistry protocols. (PDF)

